# ASOs are an effective treatment for disease-associated oligodendrocyte signatures in premanifest and symptomatic SCA3 mice

**DOI:** 10.1101/2022.07.18.500473

**Authors:** Kristen H. Schuster, Annie J. Zalon, Danielle M. DiFranco, Alexandra F. Putka, Nicholas R. Stec, Sabrina I. Jarrah, Arsal Naeem, Zaid Haque, Hanrui Zhang, Yuanfang Guan, Hayley S. McLoughlin

## Abstract

Spinocerebellar ataxia type 3 (SCA3) is the most common dominantly inherited ataxia. Currently, no preventative or disease-modifying treatments exist for this progressive neurodegenerative disorder, although efforts using gene silencing approaches are under clinical trial investigation. The disease is caused by a CAG repeat expansion in the mutant gene, *ATXN3*, producing an enlarged polyglutamine tract in the mutant protein. Similar to other paradigmatic neurodegenerative diseases, studies evaluating the pathogenic mechanism focus primarily on neuronal implications. Consequently, therapeutic interventions often overlook non-neuronal contributions to disease. Our lab recently reported that oligodendrocytes display some of the earliest and most progressive dysfunction in SCA3 mice. Evidence of disease-associated oligodendrocyte signatures has also been reported in other neurodegenerative diseases, including Alzheimer’s disease, ALS, Parkinson’s disease, and Huntington’s disease. Here, we assess the effects of anti-*ATXN3* antisense oligonucleotide (ASO) treatment on oligodendrocyte dysfunction in premanifest and symptomatic SCA3 mice. We report a severe, but modifiable, deficit in oligodendrocyte maturation caused by the toxic gain-of-function of mutant ATXN3 early in SCA3 disease that is transcriptionally, biochemically, and functionally rescued with anti-*ATXN3* ASO. Our results highlight the promising use of an ASO therapy across neurodegenerative diseases that requires glial targeting in addition to affected neuronal populations.

## INTRODUCTION

Spinocerebellar ataxia type 3 (SCA3), also known as Machado-Joseph disease, is the most common dominantly inherited ataxia in the world^1^. It is caused by a CAG trinucleotide repeat expansion in exon 10 of the *ATXN3* gene, which translates to a hyperexpanded polyglutamine (polyQ) tract in the ATXN3 protein. SCA3 patients experience progressive loss of motor control^2-4^ with neuronal death and gliosis in specific regions of the brain, including the deep cerebellar nuclei (DCN), pontine nuclei, and spinocerebellar tract^5-7^. To date, no treatments exist for this uniformly fatal disorder. Current research focusing on identifying targets for therapeutic intervention has demonstrated gene suppression to be a promising strategy, as silencing of *ATXN3* is well-tolerated in mouse models of SCA3^8-11^.

Of the available gene suppression methods, antisense oligonucleotides (ASOs) are a powerful, non-viral tool for the treatment of neurological disorders. ASOs involve a single-stranded oligonucleotide binding a complimentary target mRNA to prevent translation. Numerous ongoing preclinical and clinical trials aim to assess ASO efficacy for the treatment of neurodegenerative diseases that have historically lacked viable therapeutic options^12-16^. Toward SCA3 disease, previous proof-of-concept studies established the safety and efficacy of ASOs as a disease-modifying therapy in SCA3 mouse models^9,10,17-20^. Our group has demonstrated widespread distribution of an anti-*ATXN3* ASO (hereafter referred to as ASO-5) throughout the central nervous system (CNS) after intracerebroventricular (ICV) injection of an 8-week-old symptomatic mouse^9^. ASO-5 effectively targeted SCA3-relevant neuronal populations in areas of the brain that are particularly vulnerable to degeneration, including pontine and deep cerebellar nuclei^9,18^. Treatment with ASO-5 in homozygous SCA3 (Q84/Q84) mice decreased nuclear accumulation of ATXN3 and expression of toxic oligomeric high molecular weight ATXN3, in addition to rescuing locomotor activity after symptomatic onset^18^. A recent study demonstrated that cerebellar and brainstem neurochemical signatures in Q84/Q84 mice mirror those in patients with SCA3, and that ASO-5 treatment reversed these neurochemical abnormalities^21^. In combination, this research provides evidence that ASO mediated ATXN3 suppression is a well-tolerated therapeutic approach that rescues several disease-relevant phenotypic hallmarks in a symptomatic mouse model of SCA3.

Although ASOs have been shown to be a compelling treatment strategy in SCA3, an outstanding gap in this field is the effect of ASOs on non-neuronal cells. Most research investigating the mechanisms contributing to neurodegeneration in SCA3 has focused on neuronal dysfunction. Previous work from our lab established that among the earliest and most progressive changes in SCA3 disease is an oligodendrocyte (OL) maturation defect^22^. Through RNA sequencing (RNAseq) analysis, we identified robust differentially expressed genes implicating OL dysfunction in disease pathogenesis. With transcriptional, biochemical, and histological verification, we confirmed impairments in OL maturation that corresponded to abnormal myelination in SCA3 mice^22^. Our lab further demonstrated the spatiotemporal progression of OL maturation impairments in an SCA3 mouse model to parallel the disease progression seen in SCA3 patients^23^. This deficit has also been corroborated in SCA3 post-mortem cerebellar tissue, with significant white matter loss shown by histological and biochemical analysis^23,24^. In addition, recent reports of neurometabolite abnormalities via MRS in SCA3 mice have shown that total choline levels, indicative of OL dysfunction, are rescued with ASO-5 treatment^21^.

However, the question of whether ASO-5 therapy has an effect on the OL maturation impairment has not been addressed. Similar disease-associated OL signatures have been reported in multiple other neurodegenerative diseases, including other SCAs, Alzheimer’s disease, Amyotrophic Lateral Sclerosis (ALS), Parkinson’s disease, and Huntington’s disease^25-29^. Many of these neurodegenerative disease programs also have preclinical and clinical development of ASO therapy^12-16,29^, but have not yet explored in-depth the therapeutic efficacy in targeting and rescuing disease-associated OL signatures. Therefore, in this study we utilized SCA3 mice as a paradigmatic model of OL dysfunction in neurodegenerative diseases to test the efficacy of ASO treatment on SCA3 OL maturation impairments.

We first address whether OL maturation impairments can be prevented by treating SCA3 mice with ASO-5 prior to motor dysfunction. We show the impact of premanifest suppression of *ATXN3* on secondary phenotypes, including weight and motor activity, as well as on the onset and magnitude of OL transcriptional impairments across three vulnerable brain regions: the brainstem, cerebellum, and spinal cord. The majority of ASO treatments are being evaluated in early symptomatic patients, therefore we next assess if ASO-5 treatment in symptomatic SCA3 mice alleviates OL dysfunction. We quantify the magnitude of rescue of OL maturation transcripts and protein across the same three vulnerable brain regions in a symptomatic SCA3 mouse model. For a more in-depth understanding of the pathways that may underlie dysfunction in disease and rescue of OL maturation impairments with ASO-5 treatment, we transcriptionally profile the most robustly affected region, the brainstem. To confirm functional rescue of myelinating OLs, we assess axonal myelination in a vulnerable white matter tract via TEM analysis. By evaluating premanifest and symptomatic injection of an anti-*ATXN3* ASO, we determine the efficacy of ASO-5 treatment on OL maturation defects in a paradigmatic neurodegenerative disease and emphasize the importance of targeting non-neuronal cells in future therapeutic interventions.

## RESULTS

### PRE-SYMPTOMATIC ANTI-ATXN3 ASO DELIVERY PREVENTS LOCOMOTOR AND WEIGHT PHENOTYPES IN SCA3 MICE

SCA3 is believed to be caused primarily by the gain-of-toxic-function of mutant ATXN3. Because of this, we first sought to determine whether knocking down *ATXN3* prior to disease onset would hinder OL transcriptional impairments. We previously reported dysregulation of OL transcripts begins between 3 and 4 weeks of age in the brainstem tissue of homozygous Q84 mice^22^. Interestingly, this immediately follows the end of the highest rate of myelination in mice, which occurs between post-natal day 7 and 21^30^. To ensure delivery of ASO-5 therapy before OLs begin to mature, we treated post-natal day 0 (P0) mice via stereotactic ICV injection with 20 µg of ASO-5 (**Fig 1A**). Weights were recorded weekly for 4 weeks, after which locomotor activity was assessed and tissue was collected (**Fig 1B**). Only at 4 weeks of age did the weight of vehicle treated disease and wildtype (WT) mice statistically differ, but ASO-5 treatment prevented this weight reduction (**Fig 1C**). At 4 weeks old, ASO-5 treated disease mice were indistinguishable by weight from their WT littermates (**Fig 1C**). This is important to note, as our previously reported ASO treatment in adult-aged symptomatic mice did not have any restorative effect on weight^18^. Assessment of motor impairments at 4 weeks of age in Q84 vehicle treated mice relative to WT vehicle treated littermates demonstrated a significant reduction in both total motor activity and rearing behavior (**Fig 1D-E**). Again, ASO-5 treated disease mice were indistinguishable from WT littermates on both total locomotor activity and rearing behavior assessments at this timepoint (**Fig 1D-E**). As this was the first study to evaluate ASO treatment in neonate SCA3 mice, we established the spread of ASO expression throughout the brain. Utilizing an anti-ASO antibody, we verified consistent ASO expression confined within the CNS with no spread to peripheral tissues (**Fig 1F**). These data suggest early treatment to silence mutant *ATXN3* may be an effective strategy in delaying or preventing the onset of SCA3, highlighting the importance of identifying quantifiable biomarkers early in disease.

**Figure 1.**
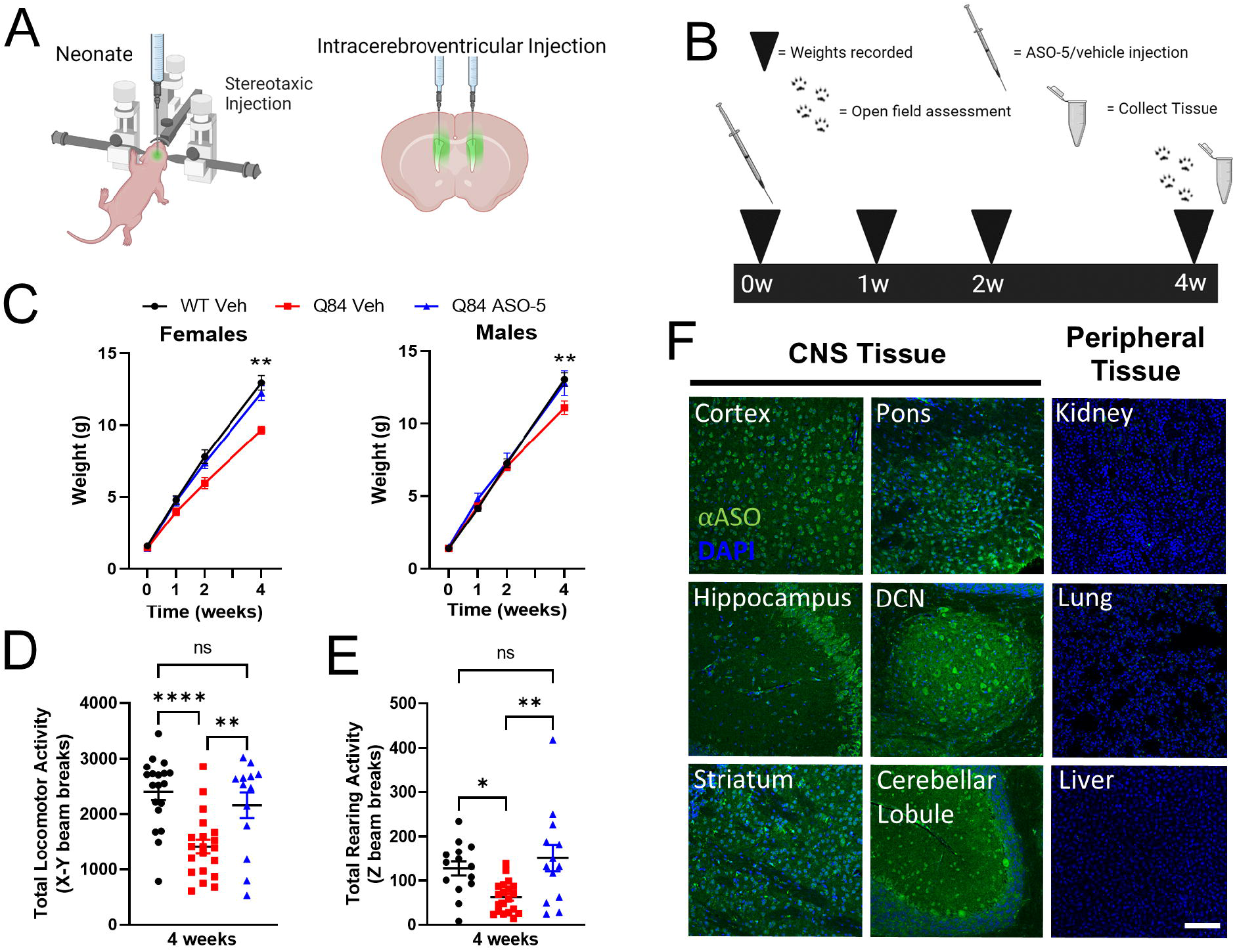
Neonate anti-*ATXN3* ASO injection prevents both weight loss and locomotion impairments despite concentrated delivery in CNS tissue. **(A)** Illustration of neonate intracerebroventricular (ICV) injections. **(B)** Schematic of longitudinal pre-symptomatic study design. Q84 and WT mice were injected with either 20 µg of ASO-5 or vehicle (PBS) treatment at post-natal day 0. Weights were recorded at 0, 1, 2, and 4 weeks of age. Open field assessment was performed prior to tissue collections at 4 weeks. **(C)** Longitudinal female and male weights from each of the 3 treatment groups. By 4-weeks-old, the weights of both male and female Q84 ASO-5 treated mice were similar to WT weights, while Q84-vehicle treated mice were significantly smaller than both other treatment groups. Open field assessment measuring **(D)** total number of X-Y beam breaks and **(E)** rearing activity of WT vehicle, Q84 vehicle, and Q84 ASO-5 treated mice at 4 weeks of age. Q84 vehicle treated mice were significantly less active than ASO-5 treated Q84 and vehicle treated WT mice, with ASO-5 treatment preventing the onset of motor and exploratory behavioral impairment at 4 weeks of age. **(F)** Representative sagittal sections of ASO-5 immunofluorescence (green) of 4-week-old ASO-5 treated SCA3 mouse brain and peripheral tissues, co-stained with DAPI (blue). ASO-5 effectively targets the whole brain, without immunofluorescence detection in peripheral tissues. Scale bar 100 µm. DCN = deep cerebellar nuclei. For C-E, data (mean ± SEM) are reported (n=5-20 mice per group). Mixed effects analysis with post-hoc Tukey’s multiple comparisons test was performed in C. One-way ANOVA with a Tukey’s post hoc analysis was performed in D and E (ns=not significant, *p<0.05, **p<0.01, ****p<0.0001).

### PREMANIFEST ASO-5 TREATMENT SUPPRESSES MUTANT ATXN3 AND PREVENTS TRANSCRIPTIONAL OLIGODENDROCYTE MATURATION IMPAIRMENTS IN VULNERABLE SCA3 BRAIN REGIONS

To confirm ASO-5 treatment in P0 mice broadly targets and silences *ATXN3* expression throughout the CNS, we transcriptionally analyzed three of the most vulnerable brain regions in SCA3, the brainstem, cerebellum, and spinal cord at 4 weeks of age. We found that ASO-5 treatment in disease mice decreases expression of both the human and mouse *ATXN3* transcript from brainstem and spinal cord tissue relative to vehicle treated disease mice (**Fig 2A-B**). Cerebellar tissue showed no significant differences in ASO-mediated *ATXN3* gene silencing (**Fig 2A-B**), however the most affected subregion within the cerebellum is the DCN, which comprises only a small part. Monomeric mutant ATXN3 (mutATXN3) was decreased significantly only in the cerebellum at 4 weeks after treatment (**Fig 2C-D**), while high molecular weight (HMW) aggregated ATXN3 protein was significantly reduced in the spinal cord samples and trending towards significantly decreased in the brainstem and cerebellum (**Fig 2C, E)**.

**Figure 2.**
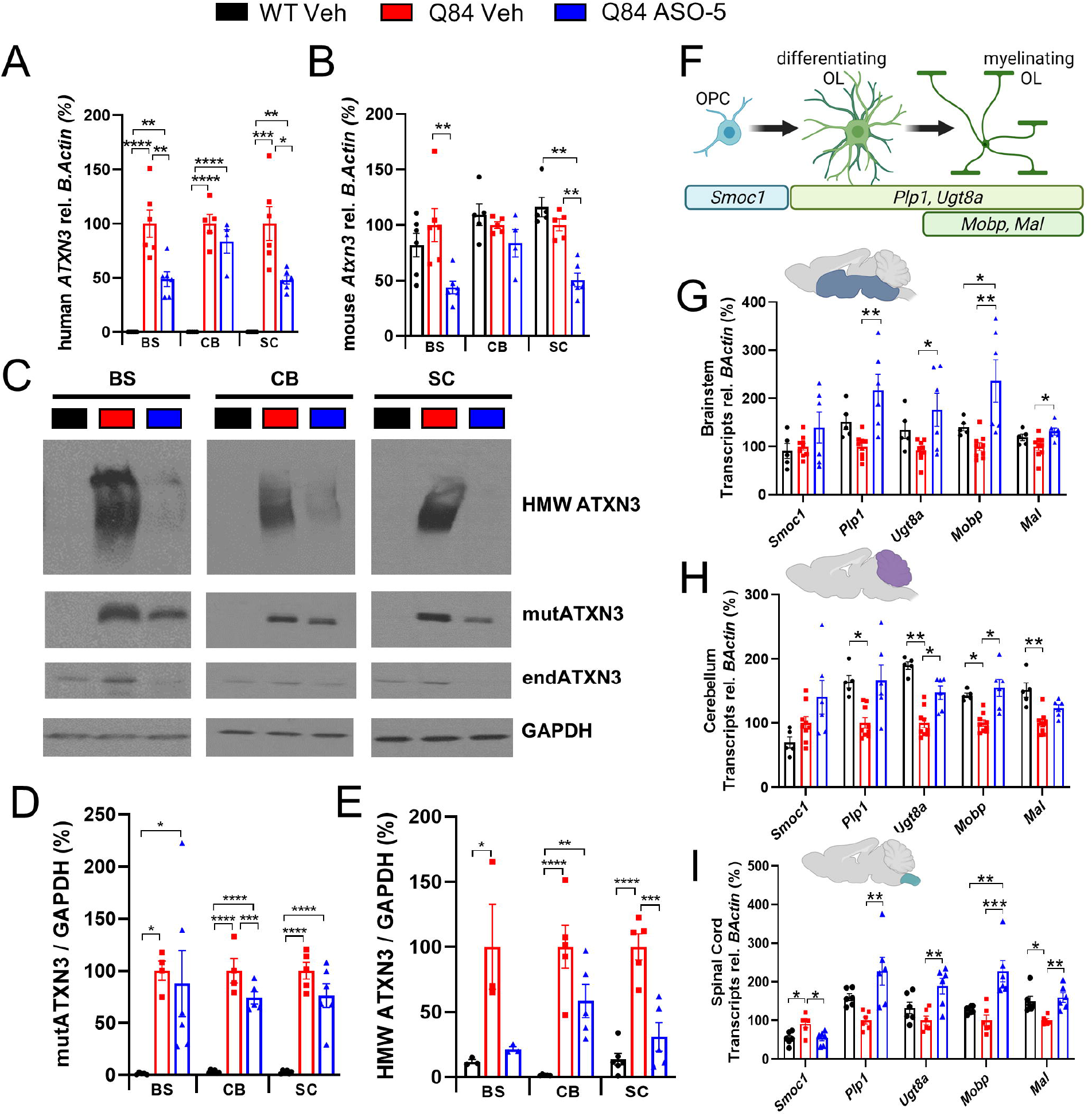
Anti-*ATXN3* ASO injections in pre-symptomatic SCA3 mice reduce ATXN3 levels and rescue OL maturation transcriptional impairments. Quantification of **(A)** human mutant *ATXN3* and **(B)** endogenous mouse *Atxn3* transcript expression in the brainstem (BS), cerebellum (CB), and cervical spinal cord (SC) in 4-week-old treated mice. ASO-5 treatment reduced human mutant *ATXN3* and endogenous mouse *Atxn3* in both brainstem and spinal cord tissue. **(C)** Representative western blot of high molecular weight ATXN3 species (HMW ATXN3), mutant ATXN3 (mutATXN3) and endogenous ATXN3 (endATXN3) protein expression across tissue types at 4 weeks of age. Quantification of **(D)** mutATXN3 and **(E)** HMW ATXN3 protein expression in the brainstem, cerebellum, and cervical spinal cord in treated mice. Q84 ASO-5 treatment relative to Q84 vehicle results in a reduction of HMW ATXN3 in all brain regions assessed. **(F)** Schematic of OL transcripts and corresponding maturation state. OL maturation transcript levels in the **(G)** brainstem, **(H)** cerebellum, and **(I)** cervical spinal cord in 4-week-old treated mice. OPC transcript *Smoc1* is increased in diseased spinal cord, while mature OL transcripts (*Plp1, Mobp, Ugt8a*, and *Mal1*) are decreased in all tissues assessed. Dysregulation of the majority of OPC and mature OL transcript was prevented with ASO-5 treatment in all brain regions assessed. Data (mean ± SEM) are reported relative to Q84 vehicle treated samples (n=3-6 mice per group). One-way ANOVA with a Tukey’s post hoc analysis was performed (no bar = not significant, *p<0.05, **p<0.01, ***p<0.001, ****p<0.0001).

Dysregulation in mature myelinating OL transcripts begins between 3 and 4 weeks of age in the brainstem region of Q84 mice^22^, therefore we addressed if the onset of this impairment could be impeded with premanifest ASO treatment throughout the mouse brain. Specifically, we previously showed normal expression of OL transcripts at 3-weeks-old, however by 4 weeks of age, there were significant increases in levels of OPC marker *Smoc1* and significant decreases in differentiating or mature oligodendrocyte markers *Ugt8a, Plp1*, and *Mobp*^22^. In ASO-treated diseased brainstem, cerebellum, and spinal cord tissues at 4 weeks of age, expression of many of these OL lineage markers were equivalent to WT levels (**Fig 2F-I**). Our previous study showed that changes in mature OL RNA expression at 4 weeks translated to reduced mature OL protein expression, though not until 8 weeks of age in Q84 mice^22^. Because of this, it is unlikely that there would be any evident changes in mature OL cell counts or white matter tract myelination at 4 weeks of age. Due to the study design, we were unable to evaluate the downstream translational effects at later timepoints. However, RNA recovery indicates that even in a premanifest state, these OL transcripts are relevant and modifiable. Further, these results suggest the onset of this OL-associated disease signature in SCA3 can be delayed with early ASO treatment.

To ensure that ASO-5 treatment is reducing the ATXN3 burden in OLs as well as neurons after premanifest injections, we histologically evaluated ATXN3 nuclear accumulation in cell types in the pons (**Fig 3A-C**) and DCN (**Fig 3D-F**). Indeed, in both brain regions we found that compared to WT vehicle treated mice, nuclear expression of ATXN3 was significantly increased in vehicle treated disease mice, but precluded from neurons (**Fig 3B, E**), and importantly from OLs (**Fig 3C, F**) with ASO-5 treatment.

**Figure 3.**
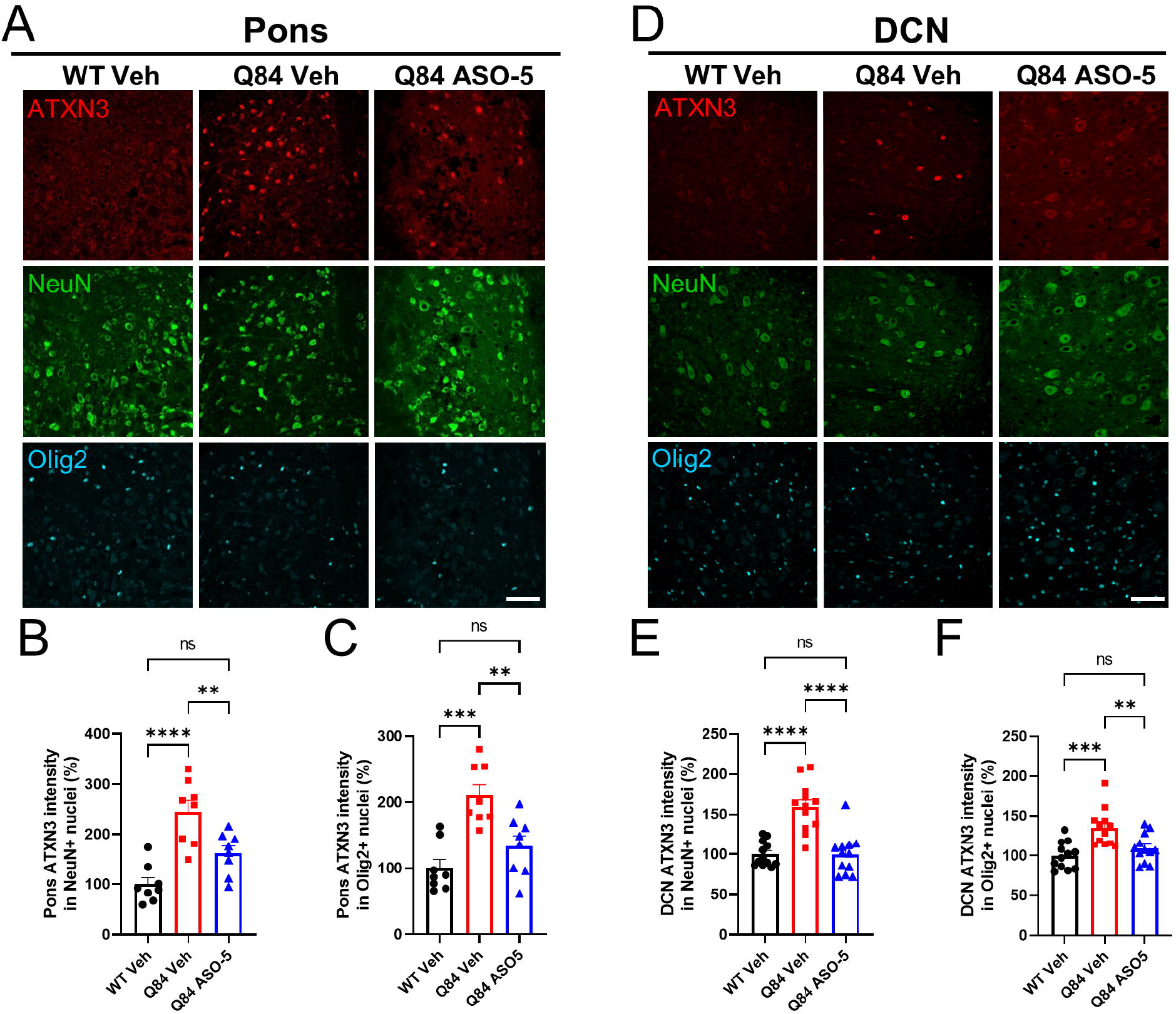
Nuclear accumulation of ATXN3 is blocked in both neurons and oligodendrocytes with pre-symptomatic ASO-5 treatment. **(A)** Representative immunofluorescent staining of ATXN3 (red), NeuN (green), and Olig2 (cyan) in the pons of WT vehicle, Q84 vehicle, and Q84 ASO-5 treated mice. Quantification of ATXN3 nuclear expression in **(B)** neurons and **(C)** oligodendrocytes. **(D)** Representative immunofluorescent staining of ATXN3 (red), NeuN (green), and Olig2 (cyan) in the DCN of WT vehicle, Q84 vehicle, and Q84 ASO-5 treated mice. Quantification of ATXN3 nuclear expression in **(E)** neurons and **(F)** oligodendrocytes. Scale bar 50 µm. Data (mean ± SEM) are reported relative to WT vehicle treated samples (n=4 mice per group). One-way ANOVA with a Tukey’s post hoc analysis was performed in B-C and E-F (ns=not significant, **p<0.01, ***p>0.001, ****p<0.0001).

Because we observed a decrease in ATXN3 nuclear accumulation in the DCN, it is possible that transcriptional assessment of whole cerebellum tissue is diluting out the effects on targeted SCA3 vulnerable regions (**Fig 2**). ASO-5 prevented nuclear accumulation in OLs in addition to neurons, therefore these results suggest that the hindering of OL maturation impairments may not be completely due to deterrence of neuronal dysfunction. This is consistent with previous reports from our lab that demonstrated OL maturation impairments to be a cell-autonomous gain of toxic function effect of mutant ATXN3^22,31^.

### ASO-5 TREATMENT OF A SYMPTOMATIC SCA3 MOUSE REDUCES ATXN3 AND RESCUES DISEASE-ASSOCIATED OLIGODENDROCYTE TRANSCRPTIONAL AND PROTEIN CHANGES

Prior studies from our lab established successful ASO-mediated ATXN3 suppression and behavioral rescue in the Q84 mouse model with treatment after symptomatic onset^18^. Therefore, we next sought to evaluate if OL maturation impairments were also rescued in treated symptomatic SCA3 mice. As several disease phenotypes, including behavior, histopathology, and OL maturation impairments, are evident by 8 weeks of age in SCA3 mice^18,22,32,33^, this age was defined as ‘symptomatic’. We treated 8-week-old Q84 mice with ICV injection of ASO-5 or vehicle and compared OL transcripts and protein to vehicle treated WT littermates (**Fig 4A**). Phenotypic rescue after ASO treatment has been shown to persist for up to 8 weeks^18^, therefore we harvested brain tissue from mice at 16 weeks of age and transcriptionally analyzed genes that are selectively enriched across OL maturation states in three of the most affected brain regions in SCA3: the brainstem (**Fig 4B**), cerebellum (**Fig 4C**), and spinal cord (**Fig 4D**).

**Figure 4.**
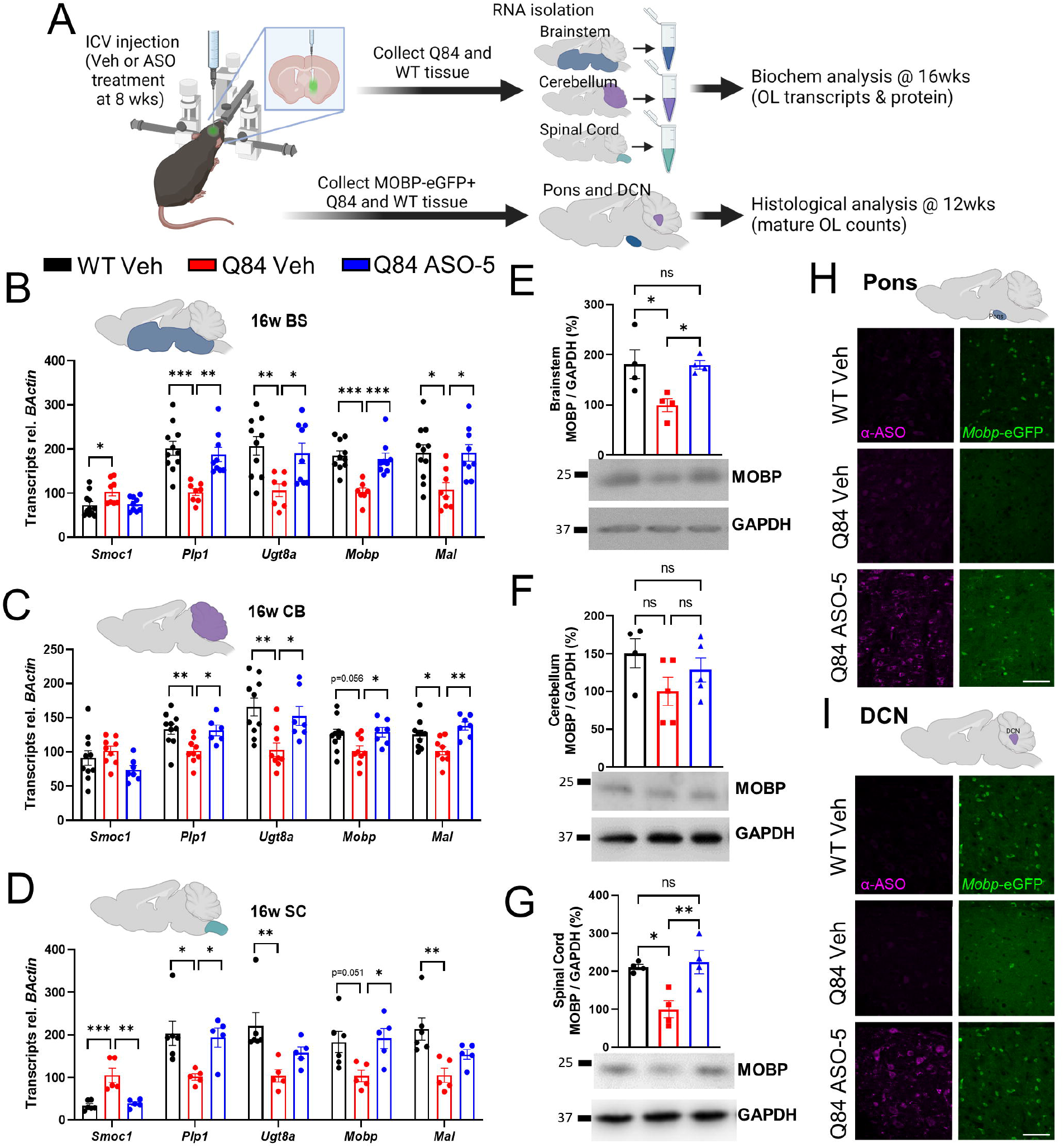
OL maturation pathway impairment is rescued in brainstem, cerebellar, and cervical spinal cord tissue with post-symptomatic ASO-5 treatment. **(A)** Schematic of ASO treatment and post-hoc biochemical and histological analysis of WT vehicle, Q84 vehicle, and Q84 ASO-5 treatment groups. OL maturation transcript levels in the **(B)** brainstem (BS), **(C)** cerebellum (CB), and **(D)** cervical spinal cord (SC) in 16-week-old mice treated 8 weeks earlier. OPC transcript *Smoc1* is increased in disease, while differentiating and mature OL transcripts (*Plp1, Ugt8a, Mobp*, and *Mal)* are decreased. The majority of transcript levels are rescued to WT levels with ASO-5 treatment in all brain regions assessed. Total protein levels of mature OL marker MOBP in the **(E)** brainstem, **(F)** cerebellum, and **(G)** cervical spinal cord. MOBP is decreased in disease brainstem and spinal cord and rescued with ASO-5 treatment. Representative sagittal **(H)** pons and **(I)** DCN images of WT and Q84 SCA3;*Mobp*-eGFP+ mice injected with ASO-5 or vehicle at 8 weeks and collected 4 weeks later (n=2-3 mice per group). *Mobp*-eGFP+ cells are reduced in Q84 vehicle treated mice and return to vehicle treated WT levels in disease mice treated with ASO-5. Scale bar 50 µm. For B-G, data (mean ± SEM) are reported relative to Q84 vehicle samples (n=3-6 mice per group). One-way ANOVA with a Tukey’s post hoc analysis was performed (no bar or ns = not significant, *p<0.05, **p<0.01, ***p<0.001, ****p<0.0001).

Assessing the same transcripts as in premanifest mice, we found OPC marker *Smoc1* to be increased in the brainstem and spinal cord, whereas markers for differentiating (*Plp1* and *Ugt8a*) and mature OLs (*Mobp* and *Mal*) in all three brain regions were decreased in disease relative to WT vehicle treated mice (**Fig 4B-D**). Consistent with premanifest treatment, symptomatic ASO-5 therapy was effective at completely restoring regional OL maturation transcripts to WT levels across all tissue samples for nearly every transcript assessed (**Fig 4B-D**). We then assessed protein changes by western blotting for MOBP, one of the three most highly expressed proteins in mature myelinating OLs. In the brainstem and spinal cord, mature OL protein changes were representative of mature OL transcript changes in that total MOBP protein levels were decreased by 50% in disease mice and rescued to 100% of WT levels with ASO-5 treatment (**Fig 4E, G**). MOBP protein levels in the cerebellum mirrored these trends, though not significantly **(Fig 4F)**. It is again important to note that in SCA3, the most vulnerable part of the cerebellum is the DCN, which comprises only a small part of the cerebellum. Therefore, although total MOBP protein levels in the cerebellum were not significantly changed, reduction and rescue of mature OL protein may be diluted by whole cerebellum analysis.

To understand the cellular implications of regional OL transcript and protein expression rescue, we examined mature OL cell counts utilizing the BAC transgenic *Mobp*-eGFP reporter mouse^34^ crossed with SCA3 Q84 (SCA3;*Mobp*-eGFP) mice. We previously used this mouse model cross to show that in SCA3 vulnerable brain regions (pons and DCN), mature *Mobp-*eGFP positive OLs are significantly reduced in an *ATXN3* dose-dependent manner^22^. As previously mentioned, this reduction was shown to be cell-autonomous and due to the gain of toxic function of mutant ATXN3^22,31^. Thus, we questioned if ASO-5 treatment could rescue this OL maturation deficit at a timepoint in which behavior is also rescued^18^. To determine if mature OL cell counts return to WT levels post-ASO-5 injection, we treated 8-week-old SCA3;*Mobp*-eGFP mice with ASO-5 and histologically evaluated the pons and DCN at 12 weeks of age (**Fig 4A**). In both brain regions, we observed a reduction of mature OLs in vehicle treated disease mice compared to vehicle treated WT mice, whereas ASO-5 treatment in disease mice rescues this phenotype (**Fig 4H-I**). Overall, these results demonstrate the efficacy of anti-*ATXN3* ASO in rescuing OL maturation impairments at the transcriptional, biochemical, and cellular level in symptomatic SCA3 mice.

### MYELINATION PATHWAY TRANSCRIPTS ARE RESCUED WITH ASO-5 TREATMENT IN AN SCA3 VULNERABLE BRAIN REGION

To achieve a greater understanding of the intracellular mechanisms that are affected by ASO-5 treatment and that may underlie the rescue of OL maturation impairments, we extracted RNA from the most robustly affected brain region in disease, the brainstem, and transcriptionally profiled each treatment group via RNAseq (**Fig 5A**). All tissue samples assessed by paired end RNAseq had an RNA integrity number (RIN) greater than seven. The average number of read counts per brainstem sample was 50.5 million (SD ± 7.5 million), with an average of 83.4% of sequencing reads mapping to exons of known genes.

**Figure 5.**
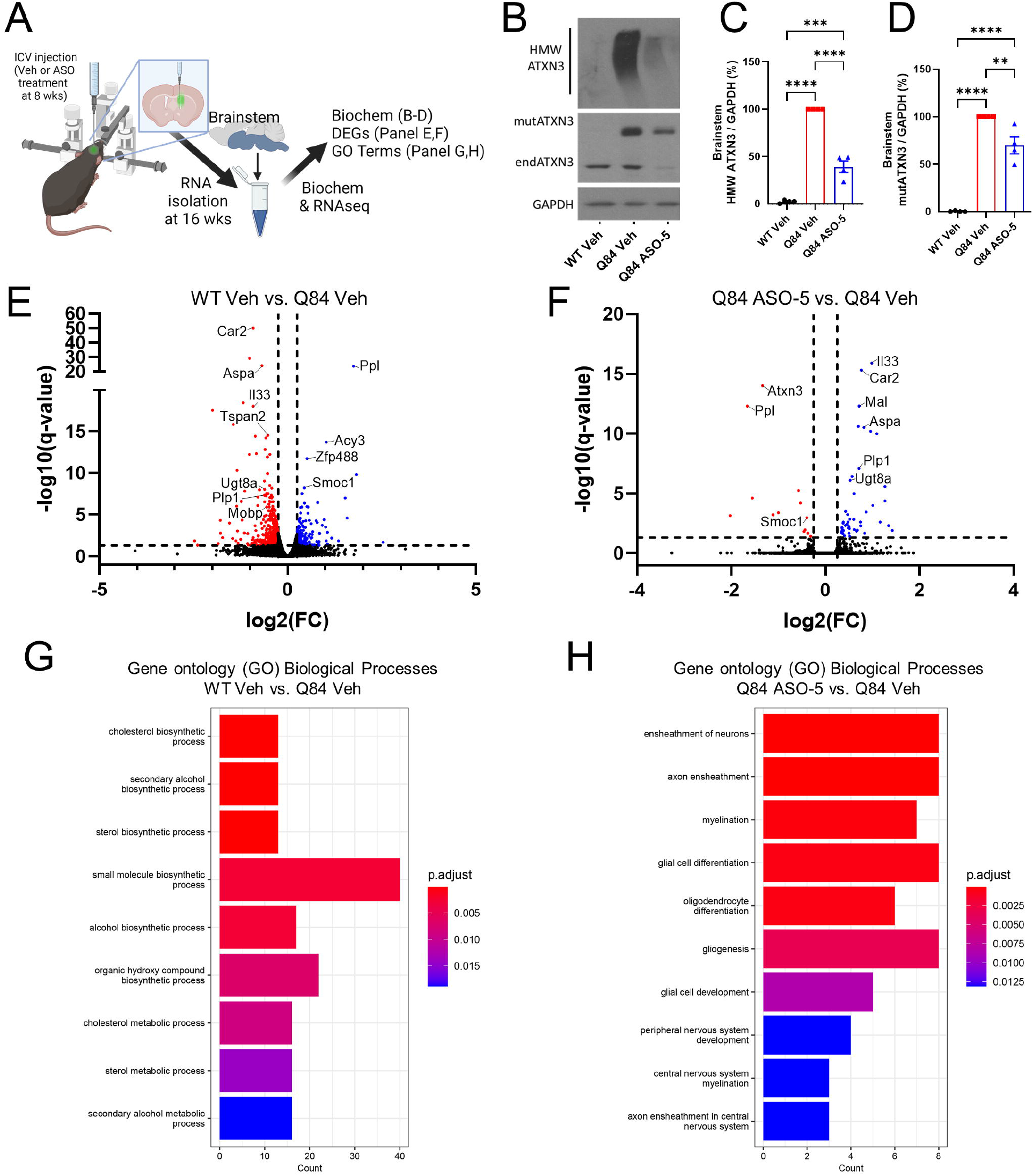
Anti-*ATXN3* ASO injections in a symptomatic SCA3 mouse reduce ATXN3 levels and rescue myelination pathways. **(A)** Overview of ASO-5 or vehicle treatment in 8-week-old SCA3 mice, isolation of brainstem tissue at 16 weeks of age, and analysis of samples. **(B)** Representative image of confirmation of the targeted partial knockdown of human ATXN3 protein by ASO-5 gene silencing in brainstem. Quantification of **(C)** HMW ATXN3 and **(D)** mutant ATXN3 protein expression in treatment groups. Volcano plots of differentially expressed genes between **(E)** WT vs Q84 vehicle treated samples or **(F)** Q84 ASO-5 treated vs Q84 vehicle treated mice. GO term analysis highlighting significant biological process between **(G)** WT and Q84 vehicle treated mice and **(H)** ASO-5 treated Q84 and vehicle treated Q84 mice in brainstem samples. n=4-5 mice per group for all experiments. For C-D, data (mean ± SEM) are reported relative to Q84 vehicle samples. One-way ANOVA with a Tukey’s post hoc analysis was performed (**p<0.01, ***p<0.001, ****p<0.0001).

RNAseq analysis confirmed suppression of human *ATXN3* gene expression in ASO treated disease mice relative to their vehicle treated counterparts in the brainstem. We validated transcript knockdown with protein analysis via Western blot and found that compared to vehicle treatment, ASO-5 treatment in disease mice partially reduced both aggregated (HMW) and monomeric (mutATXN3) disease-associated ATXN3 in brainstem tissue (**Fig 5B-D**). Bioinformatic analysis of differentially expressed (DE) gene expression (q<0.05) revealed 750 DE genes between WT and Q84 vehicle treated mice (**Fig 5E**). ASO-5 treatment resulted in only 9 DE genes relative to WT vehicle treated, however 51 DE genes were uncovered between ASO-5 and vehicle treated disease brainstem tissue (**Fig 5F**). We utilized gene ontology (GO) term analysis to determine the pathways most affected in disease and most rescued with ASO-5 treatment, as indicated by DE genes. The top biological processes disrupted in SCA3 diseased mice compared to vehicle injected WT controls were cholesterol biosynthetic, secondary alcohol biosynthetic, and sterol biosynthetic processes (**Fig 5G**). While none of these pathways directly involve OLs, cholesterol is an integral component of myelin^35,36^, suggesting some amount of myelin disruption in SCA3 brainstem tissue. Interestingly, top processes changed with ASO-5 treatment in Q84 mice predominately involved OLs, including axonal ensheathment, myelination, and OL differentiation (**Fig 5H**). This data confirms that effective therapeutic interventions for SCA3 will likely need to target OLs and myelination pathways in order to improve disease phenotypes.

### ASO-5 TREATMENT ACTIVATES MYELINATION IN BOTH VULNERABLE AND NON-VULNERABLE SCA3 BRAIN REGIONS

To assess whether rescued myelination pathway transcripts functionally restore myelin-forming capabilities of OLs, we measured axonal myelination in a brainstem white matter tract, the corticospinal tract (CST), known to be an SCA3 vulnerable brain region in both patients and mouse models ^22,37^. Again, 8-week-old control Q84 and WT mice were injected with vehicle, while treatment group Q84 mice were injected with ASO-5. Tissue was collected and fixed 8 weeks later for TEM analysis of g-ratio and axon caliber at 16 weeks of age (**Fig 6A, left bottom**). The diameter of CST axons was unchanged between vehicle treated groups while g-ratio was increased, signifying thinner myelination in Q84 controls compared to WT controls (**Fig 6A, C-D**). Importantly, ASO-5 treatment in Q84 mice completely rescued myelination, as indicated by g-ratio (**Fig 6A, C**). ASO-5 treatment also led to an increased axon diameter in the CST relative to vehicle treated mice (**Fig 6A, D**). Interestingly, the percentage of degenerated axons is not significantly different between treatment groups in this region **(Fig 6B**), suggesting that neither the impairment nor the rescue of OL maturation is due to exogenous neuronal signaling. Comparison of g-ratio to axonal caliber shows that larger axons in SCA3 vehicle treated mice have thinner myelination relative to WT vehicle treated mice, and that ASO-5 treatment also rescues this phenotype (**Fig 6E**).

**Figure 6.**
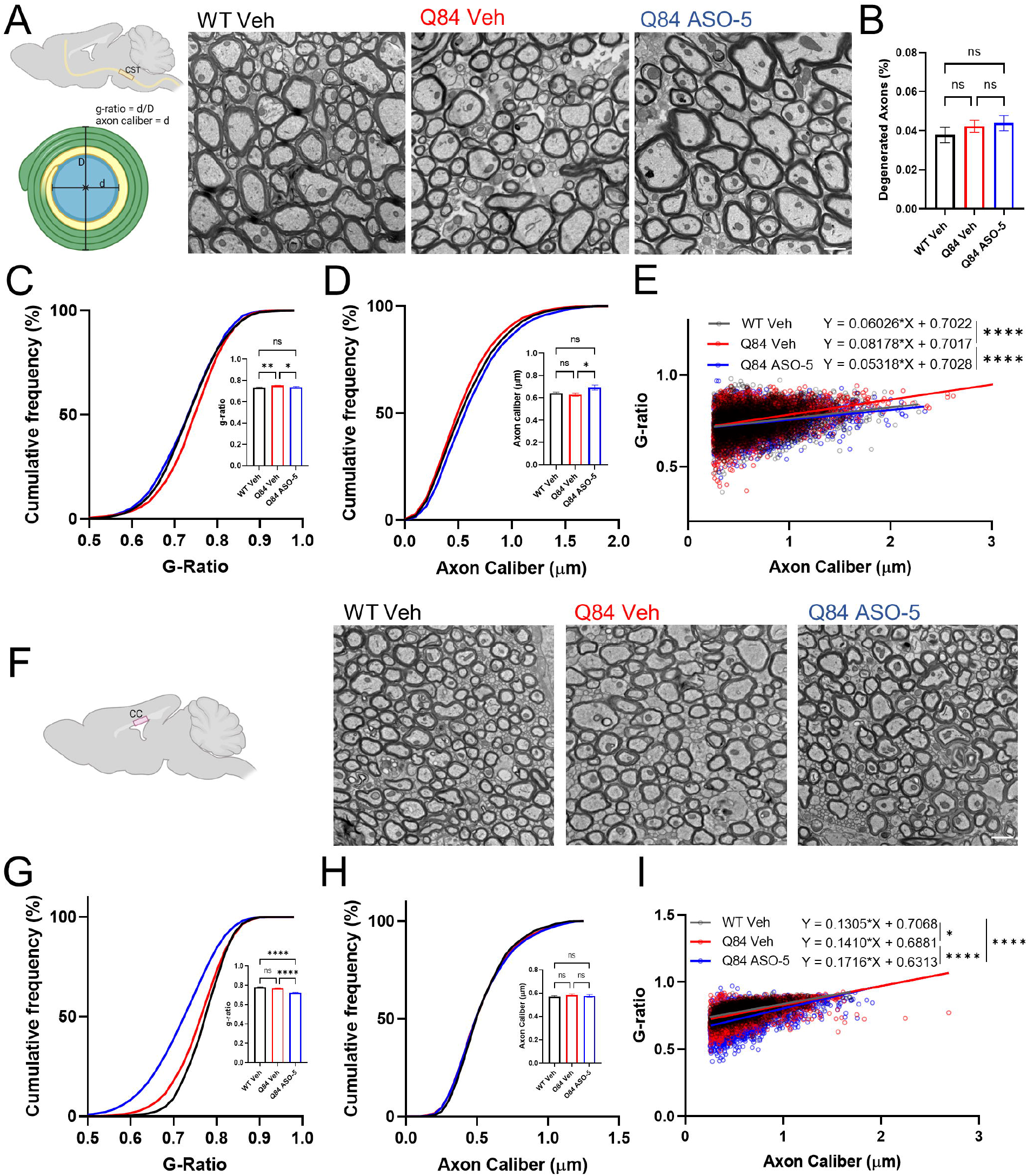
Anti-*ATXN3* ASO treatment rescues myelination in the corticospinal tract and leads to hypermyelination in the corpus callosum. **(A)** Representative coronal corticospinal tract (CST) TEM images depict abnormal myelination in 16-week-old Q84 vehicle treated mice that was rescued with ASO-5 injection 8 weeks prior. Scale bar, 1 μm. (Left bottom) Depiction of g-ratio and axon caliber measurements. **(B)** Quantification of degenerated axons in the CST show no differences between treatment groups. **(C)** Analysis of CST g-ratio indicates ASO-5 treatment rescues myelin sheath thickness to WT levels. **(D)** Analysis of CST axonal caliber shows axons in Q84 ASO-5 treated mice are significantly larger relative to Q84 vehicle treated mice, but not different from WT vehicle treated mice. **(E)** Comparison of g-ratio with axon caliber shows the consistency of myelination across axon size is rescued with ASO-5 treatment. **(F)** Representative sagittal corpus callosum (CC) TEM images. **(G)** G-ratio analyses of CC fibers reveal significantly increased myelin sheath after ASO-5 treatment in Q84 mice, while no changes are seen between Q84 and WT vehicle treated mice. **(H)** Quantification of axon diameter reveals no differences between treatment groups. **(I)** G-ratio vs. axon caliber comparison shows ASO-5 treatment affects the relationship between axon size and myelin thickness compared to vehicle treated mice. Axon caliber and g-ratio calculated using MyelTracer software (n=850-1250 axons across 10-15 images per mouse, n=3-4 mice per genotype). Data report as mean ± SEM. Simple linear regression and one-way ANOVA with a Tukey’s post hoc analysis were performed (ns=not significant, *p <0.05, **p <0.01, ****p<0.0001).

Because ASO treatment is broadly distributed throughout the brain, we were able to explore the effects of ASO-5 treatment on less vulnerable brain regions. The corpus callosum (CC) is a prominent white matter tract that we found to not have reduced mature OL cell counts at 16-weeks-old in SCA3 mice^22^. Therefore, we assessed regional OL maturation dysfunction by analyzing the ultrastructural features of myelination in this non-vulnerable white matter tract in vehicle and ASO-5 treated SCA3 mice (**Fig 6F**). As expected, we found no differences between WT and Q84 vehicle treated mice in myelination in the CC (**Fig 6G**). However, unlike ASO-5 effects in the CST, axon caliber in the CC was not changed in any of the conditions assessed (**Fig 6H**). ASO-5 treatment in SCA3 mice did not further increase the axon caliber in the CC, however it did lead to substantial increases in myelin thickness as defined by the g-ratio (**Fig 6G**), particularly in smaller sized axons (**Fig 6I**). This suggests there may be a positive-reinforcement mechanism regulating myelination that garners further mechanistic exploration. These data illustrate the complete functional and ultrastructural rescue of OLs in SCA3 and highlight the importance of considering the effects of broad-target therapies in less-affected brain regions in disease.

## DISCUSSION

Previous work from our lab established an impairment in the OL maturation process as an early and progressive feature of SCA3 pathogenesis. Disease-associated OL signatures have also recently been implicated more common neurodegenerative diseases like Alzheimer’s disease, Parkinson’s disease, ALS, and Huntington’s disease^25-28^. Many of these neurodegenerative disease programs also have preclinical and clinical development of ASO therapy that have not yet assessed the effect of treatment on oligodendrocyte signatures^12-16^. Therefore, we tested the efficacy of ASO treatment in SCA3 mice as a paradigmatic model of OL dysfunction in neurodegenerative diseases. The aim of the present study was to determine if anti-*ATXN3* ASO therapy can effectively target OLs to prevent or rescue the maturation defect in SCA3 with premanifest and symptomatic treatment, respectively. We provide evidence that ASO-5 treatment prior to symptom onset not only precludes OL transcript and protein dysregulation, but also prevents secondary phenotypes, including motor defects and weight loss. This suggests early treatment could be protective against SCA3 symptomatic onset, highlighting the need for early biomarkers of disease. Importantly, symptomatic treatment rescued OL transcriptional dysregulation in SCA3 vulnerable brain regions, which translated to both protein and ultrastructural rescue. Moreover, RNAseq analysis revealed OL maturation pathways to be pivotal to ASO-5 recovery of disease pathology. These results emphasize the importance of OL signatures in disease progression and as targets in future neurodegenerative disease therapies.

ASOs are known to broadly target all cell types of the brain – neurons and glia alike^38-41^. However, analysis of GO biological processes within brainstem tissue of ASO-5 treated Q84 mice relative to vehicle treated Q84 mice indicated heavy involvement of OLs over all other cell types: ensheathment of neurons and axons, myelination, and glial cell differentiation. Intriguingly, although direct disruption of OL maturation pathways was not seen in disease brainstem tissue compared to WT controls, the top pathway dysregulated between vehicle-treated WT and Q84 mice was cholesterol biosynthesis. Because cholesterol is a primary component of myelin sheaths^42,43^, downregulation of cholesterol biosynthesis could underlie the reduced axonal myelination seen in SCA3 CST (**Fig 6A-E**). Our findings suggest that ASO-5 confers a therapeutic effect primarily by influencing pathways related to OLs, implying that OLs could be a crucial therapeutic target in SCA3. Consistent with these results, a recent study in drosophila showed that hyperexpanded polyQ tracts in ATXN3 expressed in glia alone resulted in developmental lethality^44^, suggesting SCA3-induced glial dysfunction is sufficient to cause devastating disease phenotypes. Future investigations will need to further explore the necessity and sufficiency of both the expression and silencing of mutant ATXN3 in OLs for SCA3 pathogenesis and treatment.

A key feature among many neurodegenerative diseases is regional vulnerability. As we previously showed^22,23^ and extended here, OL maturation impairments are central to the region-specific vulnerability of nervous system pathogenesis in SCA3. The brainstem and cerebellum have been established as two of the most vulnerable brain regions in SCA3 in both patients and animal models^7^. In agreement with recent SCA3 patient literature, our findings indicate the spinal cord is also a vulnerable CNS region in SCA3 mice. Although there is little SCA3 research on animal spinal cord tissue, we found it to be as affected as the brainstem, another myelin-rich white matter brain region. Interestingly, while many white matter tracts are affected in SCA3, the CC remained unaffected. We have previously shown that this region has no change in mature OL cell counts^22^ and in this study further established the CC as a non-vulnerable brain region in SCA3 with ultrastructural analysis via TEM. Although we found no differences in myelin thickness between vehicle treated groups, ASO-5 treatment, which rescued myelination in a vulnerable brain region, surprisingly led to hypermyelination in the CC. Instead of regional vulnerability, these findings point to the possibility of regional *protection*. Much of the current work on SCA3 focuses on what causes dysfunction in vulnerable brain regions. With this study we propose a shift in the viewpoint, indicating that just as relevant are investigations of what prevents dysfunction in non-vulnerable, or protected, brain regions.

This study emphasizes the importance of considering non-neuronal contributions to disease when evaluating therapeutic interventions. OLs in particular have emerged as a key player in not only SCA3 pathogenesis, but in several other neurodegenerative diseases as well^45^, and here, we establish their therapeutic significance. While ASOs continue to be a powerful tool for SCA3 treatment, future studies into their role in rescuing dysregulated processes and possibly over-activating protective processes, in addition to their mechanism of targeting OLs, will be essential. These mechanistic studies will offer insight into the pathophysiology and therapeutic options for SCA3, as well as other neurodegenerative diseases that exhibit disease-associated oligodendrocyte signatures.

## MATERIALS AND METHODS

### Animals

The University of Michigan Institutional Animal Care and Use Committee (IACUC) approved all animal procedures. They were conducted ethically in accordance with the U.S. Public Health Service’s Policy on Human Care and Use of Laboratory Animals. All animals were housed in a room following standard 12-hour light/dark cycles with food and water provided ab libitum.

In this study, homozygous YACMJD84.2Q-C57BL/6 transgenic mice (referred to in this study as “Q84 mice”) and wild type (WT) sex- and age-matched controls were used. Genotyping was completed via tail biopsy DNA isolation prior to weaning and confirmed postmortem. The number of CAG repeats in the human *ATXN3* gene in Q84 mouse DNA was determined by fragmentation analysis (Laragen, San Diego, CA) as previously described^22^ with all enrolled mice having a minimum CAG repeat average greater than 75. Genotyping for *Mobp*-eGFP mice was completed as previously described^22^. For all studies, 4-6 mice (equal numbers of males and females) per genotype were collected for tissue analysis at 4, 12, or 16 weeks of age. Left hemisphere brain tissue was macrodissected into brainstem (includes whole of brainstem, stratum, and thalamus), cerebellum, and cervical spinal cord for biochemical analysis as previously described^9^. PBS perfused right hemispheres were post-fixed in 4% paraformaldehyde and embedded in 30% sucrose for histological assessments.

### Antisense Oligonucleotides

ASO-5 used in this study targets the human and mouse ATXN3 3’-UTR (ASO-5; GCATCTTTTCATTACTGGC). ASO-5 gapmer design and synthesis were previously described^9^.

### Surgical Delivery of Antisense Oligonucleotides

Premanifest mice used in this study received a post-natal day 0 injection of either ASO-5 or PBS into both lateral ventricles as described previously^46^. Pups were placed on ice for cryo-anesthetization before performing a bi-lateral intracerebroventricular (ICV) injection of either ASO-5 or PBS (2 µL per hemisphere, 5 µg/µL ASO-5)^46^. Immediately following injection, pups were placed on a heating pad before returning to their cage. Pups were monitored and weighed weekly following treatment. Mice were harvested at 4 weeks of age for biochemical and histological assessment as previously described^9,47^.

Symptomatic ASO treatment was completed as previously described^9,47^. Post-operative care was performed for at least 7 days following treatment.

### RNA Isolation and Sequencing

RNA was extracted from PBS-perfused, flash-frozen brainstem, spinal cord, and cerebellum samples from the left hemisphere. Using the Next Advance Bullet Blender, samples were homogenized in radioimmunoprecipitation assay (RIPA, by Sigma, St. Louis, MO) buffer (750 µL for brainstem and 500 µL for cerebellum) with added proteinase inhibitor (BioRad, Hercules, CA). A 250 µL aliquot of this lysate was reserved for Western blot analysis and the remaining sample was used for RNA extraction.

RNA extraction was conducted using the QIAshredder (Qiagen Hilden, Germany), RNeasy Mini Kit (Qiagen, Hilden, Germany), and DNAse I kit (Qiagen, Hilden, Germany), then eluted in RNase-free water, according to manufacturer’s instructions. RNA samples were assessed by nanodrop to determine concentration and sample purity. For library preparation, 1 μg of total brainstem RNA was submitted to the University of Michigan Sequencing Core for Illumina HiSeq library preparation, quality control, and RNA sequencing. WT and Q84 RNA samples were chosen for RNA sequencing based on RIN number (>7 RIN) and, if applicable, CAG repeat expansion size (>average 72Q) as determined by Laragen, Inc. (Culver City, CA).

### RNA-seq Expression Analyses

The University of Michigan Bioinformatics Core generated libraries (Illumina TruSeq) from purified brain RNA samples, completed quality control analysis, and ran 75×75 paired end sequencing on Illumina HiSeq 4000. The RNAseq data were processed as previously described^22^. Briefly, reads were pseudo-aligned to the ENSEMBL 108 mouse reference cDNA sequences (June 2020, GRCm39) using Kallisto 0.45 and quantified transcripts aggregated by their gene were fed to Sleuth 0.30.1 for differential expression analysis. Genes with expression levels less than 5 transcripts per million (TPM) in more than 25% of the analyzed samples were discarded from the differential expression analysis. We used Sleuth to evaluate statistical significance of the differential gene expression in each of the three treatment groups:

WT vehicle treated, Q84 vehicle treated, and Q84 ASO-5 treated. Through Benjamini-Hochberg correction, genes with q-values less than 0.05 and |log2FC| > 0.25 were defined as differentially expressed. The differentially expressed genes were then used for Gene Ontology (GO) term analysis by clusterProfiler 4.4.4 comparing the biological processes that were impaired in SCA3 compared to WT mice and that were resolved with ASO treatment in disease mice.

### QPCR

Quantitative PCR (qPCR) analysis was performed by reverse transcribing 0.5-1 μg of total RNA using the iScript cDNA synthesis kit according to the manufacturer’s instructions (Bio-Rad, Hercules, CA). iQ quantitative PCR was performed on the diluted cDNA following the manufacturer’s protocol (Bio-Rad, Hercules, CA) using the following primers: human *ATXN3* (Forward primer: CCTCAATTGCACATCAGCTGGAT; Reverse primer: AACGTGCGATAATCTTCACTAGTAACTC; Probe Seq: CTGCCATTCTCATCCTC), mouse *Atxn3* (Thermofisher; Cat # Mm01336273), *Beta Actin* (Thermofisher; Cat # Mm02619580), *Smoc1* (Thermofisher; Cat #Mm00491564), *Plp1* (Thermofisher; Cat #Mm01297210), *Ugt8a* (Thermofisher; Cat #Mm00495930), *Mobp* (Thermofisher; Cat #Mm02745649), and *Mal* (Thermofisher; Cat #Mm01339780). Gene expression fold change (RQ) was calculated relative to vehicle treated Q84 samples for each timepoint and normalized to *beta actin*, where RQ = 2-(ddCT).

### Open Field Motor Evaluation

Motor assessments of Q84 mice were conducted when mice were 4 weeks old. A photobeam open-field apparatus (San Diego Instruments, San Diego, CA) was used in order to measure total locomotor activity (x/y-axis beam breaks), and exploratory behavior (z-axis beam breaks). Trials lasted for 30 minutes. Experimenters were blinded to genotypes and treatment groups during behavioral tests.

### Western Blot

Following brainstem, cerebellum, and spinal cord macrodissection, protein lysates were processed as previously described^9^ and stored at -80C. Briefly, homogenized lysates were diluted, and 20-30 µg of total lysates were resolved in 4-20% gradient sodium dodecyl sulfate-polyacrylamide electrophoresis gels. Protein was transferred to 0.45 µm nitrocellulose membranes prior to overnight incubation at 4°C with the following primary antibodies: mouse anti-ATXN3 (1H9; 1:1000, MAB5360; Millipore, Billerica, MA), mouse anti□GAPDH (1:10000, MAB374; Millipore, Billerica, MA), and rabbit anti-MOBP (1:100-1:1000; ab91405; Abcam). Following incubation, molecular weight bands were visualized by peroxidase-conjugated anti-mouse or anti-rabbit secondary antibody (1:5000; Jackson ImmunoResearch Laboratories, West Grove, PA). ECL-plus (Western Lighting; PerkinElmer, Waltham, MA) treated membranes were visualized via exposure on G:Box Chemi XRQ system (Syngene) and band intensities were quantified using ImageJ analysis software (NIH). Two gels were run per brain region and samples were normalized to Q84 vehicle treated samples within each blot.

### Immunohistochemistry

The PBS-perfused right brain hemispheres were post-fixed for 24 hours in 4% paraformaldehyde (PFA) then sucrose-embedded. Prepared brains were sectioned and stained as previously reported^22^. Briefly, sections were washed in PBS, blocked using reagents from M.O.M. kits according to manufacturer’s protocol (BMK2202, Vector Laboratories, Newark, CA), and incubated in primary antibody at 4°C overnight. Primary antibodies assessed include rabbit anti-ASO (1:5000; Ionis Pharmaceuticals, Carlsbad, CA), mouse anti-ATXN3 (1H9; 1:500, MAB5360; Millipore, Billerica, MA), and rabbit anti-Olig2 (1:500, P21954, ThermoFisher, Waltham, MA). The following day sections were washed, incubated in secondary antibody or rabbit anti-NeuN (AlexaFluor 488 conjugate, 1:100, ABN78A4, Millipore, Billerica, MA), and DAPI (Sigma, St. Louis, MO), washed, and mounted with Prolong Gold Antifade Reagent (Invitrogen, Waltham, MA). Fluorescent secondary antibodies used include AlexaFluor 488 goat anti-rabbit IgG (1:1000, A11008, Invitrogen), AlexaFluor 568 goat anti-mouse IgG (1:1000, A11004, Invitrogen) and AlexaFluor 647 goat anti-rabbit IgG (1:1000, A21245, Invitrogen). Imaging was performed using a Nikon-A1 Standard Sensitivity confocal microscope with NIS-Elements software in peripheral tissue including the kidney, lung, and liver, as well as various brain regions, including the cortex, basilar pontine nuclei (denoted as pons), hippocampus, deep cerebellar nuclei (DCN), striatum, and cerebellar lobules. CellProfiler software^48^ was used to analyze histological images. Nuclear ATXN3 protein expression was measured using fluorescence intensity within NeuN+ or Olig2+ DAPI stained nuclei.

### Transmission Electron Microscopy

Mice (1-2 males and 2 females per treatment group per region) were transcardially perfused first with 1x PBS, then with 2.5% glutaraldehyde and 4% PFA in 1x PBS. Brains were removed and postfixed overnight at 4°C in the same buffer before being transferred to 0.1M phosphate buffer, pH 7.4 for long term storage at 4°C. Processing of TEM tissue was done by University of Michigan’s Microscopy Core according to their protocol and as previously described^22^. Imaging was completed on a JEOL JEM-1400 plus transmission electron microscope (JEOL USA Inc., Peabody, MA) at an accelerating voltage of 60 kV, using either an AMT 4-megapixel XR401 or a 12-megapixel Nanosprint CMOS camera (Advanced Microscopy Techniques, Woburn, MA). MyelTracer software^49^ was used for g-ratio and axon caliber measurements. Manual counts of degenerated axons were taken as a percentage of the number of total axons used in g-ratio and axon caliber analysis per image. Experimenters were blinded to genotypes and treatment groups during TEM image analysis.

### Statistics

Prism 8.0 (GraphPad Software, La Jolla, CA) was used to perform all statistical analysis. Statistical significance was evaluated using a one-way ANOVA with a post hoc Tukey’s multiple comparisons test, mixed effects analysis with post-hoc Tukey’s multiple comparisons test, or simple linear regression. All one-way ANOVA tests assumed Gaussian distribution of residuals and equal standard deviation of each condition. Variability about the mean is expressed as mean□±□standard error of the mean (SEM). All tests set the level of significance at p<0.05.

## Data Availability

RNA sequencing data that support the findings of this study can be accessed under GEO accession # (#######). All other data supporting the findings of this study are either available within the article or are available from the corresponding author on reasonable request.

## Acknowledgements

This work was supported in part by National Ataxia Foundation SCA Young Investigator Award to H.S.M.; National Institutes of Health Grant R01-NS122751 to H.S.M., and Grant U01-NS106670 to H.S.M. We acknowledge Ionis Pharmaceuticals, who identified and generated the anti-*ATXN3* ASOs used this study. We acknowledge S. Meshinchi and D. Leroux in the Michigan Medicine Microscopy Core for TEM training and processing of TEM samples. Figure schematics were created using BioRender.com.

## Author contributions

K.H.S., A.J.Z., H.Z., Y.G., and H.S.M. designed research; K.H.S., A.J.Z., D.M.D., A.F.P., N.R.S., S.J., and H.S.M. performed research; K.H.S., A.J.Z., H.Z., D.M.D., A.F.P., N.R.S., S.J., A.N., Z.H., H.Z., Y.G., and H.S.M. analyzed data; K.H.S., A.J.Z., D.M.D., A.F.P., N.R.S., H.Z., Y.G., and H.S.M. edited the paper; K.H.S., A.F.P., and H.S.M. wrote the paper; K.H.S., A.F.P., and H.S.M. wrote the first draft of the paper; H.S.M. contributed unpublished reagents/analytic tools.

